# Pipeline for generating stable large genomic deletions in zebrafish, from small domains to whole gene excisions

**DOI:** 10.1101/2021.01.27.428552

**Authors:** Alisha Tromp, Kate Robinson, Thomas E. Hall, Bryan Mowry, Jean Giacomotto

**Author notes:** Contributed equally to this study (Alisha Tromp, Kate Robinson).

## Abstract

Here we describe a short feasibility study and methodological framework for the production of stable, CRISPR/Cas9-based, large genomic deletions in zebrafish, ranging from several base pairs (bp) to hundreds of kilobases (kb). Using a cocktail of four sgRNAs targeting a single genomic region mixed with a marker-sgRNA against the pigmentation gene *tyrosinase* (*tyr*), we demonstrate that one can easily and accurately excise genomic regions such as promoters, protein domains, specific exons or whole genes. We exemplify this technique with a complex gene family, neurexins, composed of three duplicated genes with multiple promoters and intricate splicing processes leading to thousands of isoforms. We precisely deleted small regions such as their transmembrane domains (150bp deletion in average) to their entire genomic locus (300kb deletion for *nrxn1A* for instance). We find that both the concentration and ratio of Cas9/sgRNAs are critical for the successful generation of these large deletions and, interestingly, that their transmission frequency does not decrease with increasing distance between sgRNA target sites. Considering the growing reports and debate about genetically compensated small indel mutants, the use of large-deletion approaches is likely to be widely adopted in studies of gene function. This strategy will also be key to the study of non-coding genomic regions.

## INTRODUCTION

CRISPR/Cas9 has recently revolutionised genetics and the way we can manipulate gene function^1^. Commonly, to disrupt a gene using this technology, one employs a single guide/site strategy to promote targeted small indels (addition or deletion of 1 to 20bp) inducing a frameshift and the generation of a premature stop codon. Although this strategy is straightforward and very efficient, there is growing literature reporting genetic compensatory mechanisms, *i*.*e*. the mutation is rescued/compensated by diverse mechanisms that are not yet fully understood (for details see^2-7^). Recently, we began to study the neurexin gene family in the zebrafish^8^. We experienced some challenges knocking out these genes using this traditional approach. Despite successfully generating small indels triggering a premature stop in the gene-ORFs, we were still detecting expression of full-length proteins, suggesting exon skipping or use of alternative/cryptic splicing sites in all our mutants. When one has a close look at this gene family, it seems evident that these are perfect candidates for escaping traditionally engineered mutations. Indeed, these genes, very well conserved across species, display an extreme transcriptomic complexity that makes them very prone to genetic compensatory mechanisms^9^. First, each gene presents two distinctive promoters driving two major isoforms (α and β, **figure 1A**), and most exons of each form could be alternatively spliced, leading to the expression of thousands of isoforms^9^. Second, all vertebrates present at least 3 neurexin genes (duplicated in zebrafish) with high sequence identity, making them prone to cis-regulation via non-sense mediated decay of one of the homologues (*i*.*e*. premature stop in *NRXN1* may trigger non-sense mediated decay and up-regulation of *NRXN2* and/or *NRXN3*). To avoid, but also study, genetic compensation of these genes, we endeavoured to establish mutants presenting i) a full deletion of each entire gene, ii) a specific deletion of the long α-isoforms and iii) deletion of shared domains such as the transmembrane region, essential for anchoring those proteins to the synaptic membrane. Considering the potential difficulty of these tasks (three large genes duplicated in zebrafish – six genes in total) and the absence of technical feasibility for removing hundreds of kilobases (kb), we first validated the possibility of generating such large deletions in the zebrafish.

**Figure 01:**
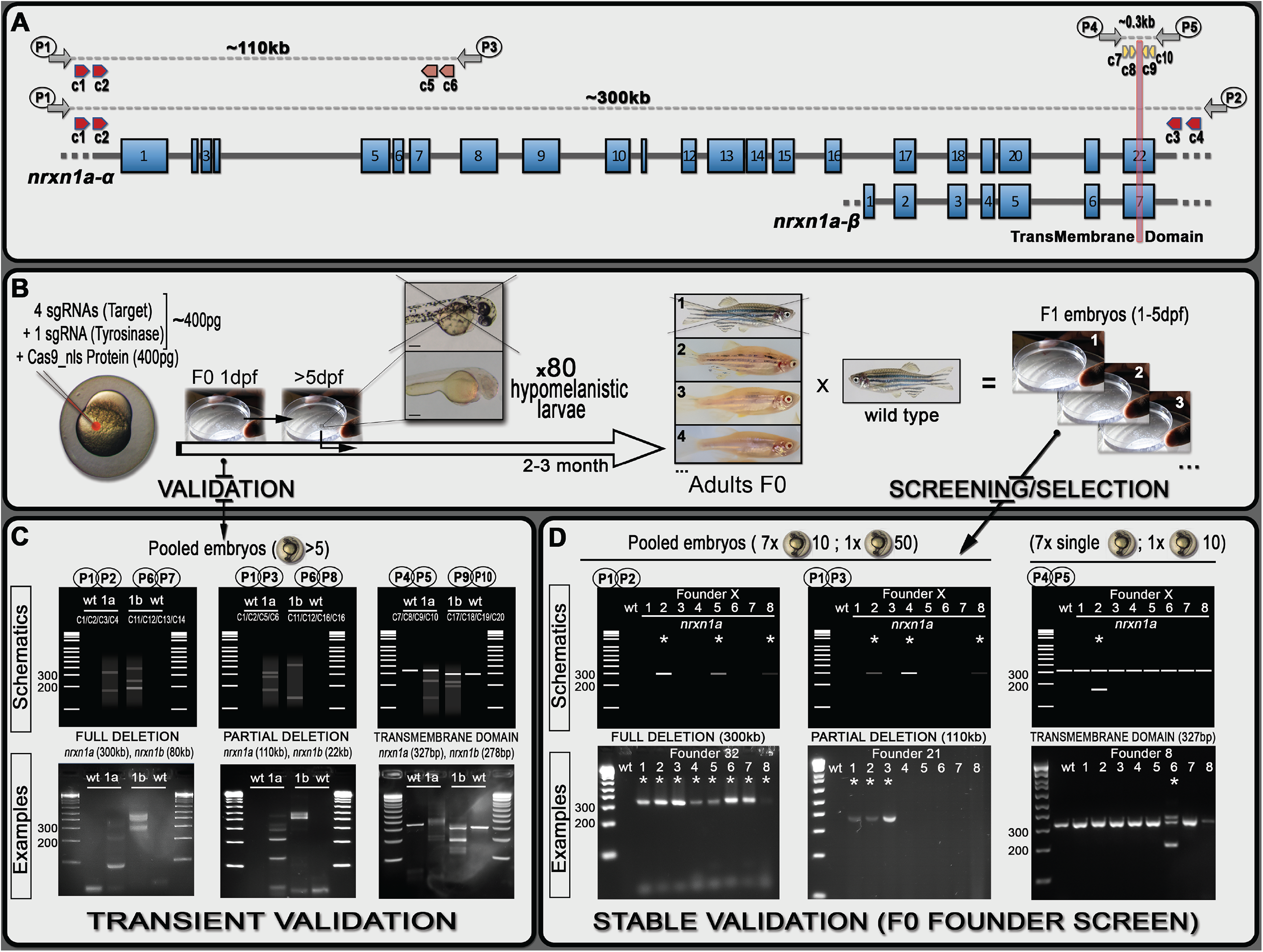
Schematics of neurexin organisation and pipeline for generating mutants with large deletions. **A**, Representation of neurexin1a (*nrxn1a*) genomic region together with sgRNA (CRISPR) target sites (c1 to c10) as well as primers used for validation and screening. The size of exons, intron gaps and protein domains (and compared to one another) are not drawn to scale. **B**, Pipeline designed to generate large deletions in zebrafish. 24 hours post-injection, pools of five embryos are collected to evaluate the efficiency of the Cas9/sgRNAs mixes (see C. Transient Validation). Following validation, 80x 5dpf-larvae are selected based on their pigmentation phenotype. Once adult, F0 animals are crossed against wildtype to identify carriers/founders able to transmit the designed deletion to F1 mutants (See D. Stable Validation). **C**, To analyse/validate the efficiency of our sgRNAs-mixes, we proceed with a basic PCR approach. Depending on the size of the target, a wildtype amplicon will be amplified or not; with the presence of a wildtype product hampering the amplification/detection of DNA remodelling (especially if rare). If large DNA deletion(s) happens, one should observe smearing with the presence of clear amplicon(s), as presented in the schematics. The lower panels present the results we obtained for *nrxn1a* and *nrxn1b* transient validations. These transient validations could also be performed on individual embryos. **D**, The same PCR approach is conducted to identify F0 adult founders. This time, 8 batches are collected as schematically described. As opposed to the transient approach, a sharp band(s) should be observed, evidencing the presence of a large deletion. Upper panels represent theoretical results for one specific founder, while lower panels present PCR profiles obtained during our screen against the different *nrxn1a* deletion design. Founder frequency was evaluated at approximately 1/10 without significant impact of the target/deletion size.

Here, we not only report that it is feasible to accurately delete large genomic regions (ranging from several kb to potentially more than 1Mb), but we also describe a pipeline to ease the selection of F0-founders **(figure 1)**. Such an approach has the potential to help generate mutants to better target/study small gene regions such as binding domains, catalytic sites, localisation signals and/or specific exons. The ability to accurately generate large deletions will also be very useful to study non-coding regions as well as enhancers and promoters. Finally, the generation of such mutants should also help the study of genes that are prone to genetic compensation as well as study those mechanisms *per se*.

## RESULTS AND DISCUSSION

### Validation of large deletion events in injected embryos

To test our ability to generate stable mutants with large genomic deletions, we initially designed a strategy based on four sgRNAs with two guides on each side of the sequence to be removed (schematic representation with *nrxn1a* gene in **figure 1A**). In theory, several steps could be performed to validate the efficiency of each individual sgRNA, such as *in vitro*, HRMA and/or T7E1 assays^10-12^. However, we demonstrate here that a simple PCR approach on a subset of the injected embryos is sufficient to visualise/validate proper deletion events. We initially injected four guides against different regions of *nrxn1a* and *nrxn1b*, aiming to delete: 1) the full genomic region (≈300kb and ≈80kb respectively), 2) the α-isoforms (≈100kb and ≈20kb respectively) and 3) the transmembrane domain (≈0.15kb) (**figure 1A, table S01**). We collected a mix of five embryos from each injection, extracted genomic DNA and performed PCR amplification with respective primers as presented in **figure 1** and **table S02**. For full and isoform-specific deletions, although no wildtype amplicon could be detected (wt-target too large to amplify), different bands as well as clear smears were noticeable with the injected embryos of each targeted gene, demonstrating DNA remodelling and the presence of large deletions in at least some cells of the collected embryos (**figure 1B-C**). For the transmembrane domain, the picture is not as evident considering that a wildtype band could be amplified, thereby reducing the relevance of a PCR/gel approach to highlight non-abundant DNA remodelling/deletions. Nonetheless, we were still able to detect smears and/or smaller bands than the wildtype amplicons in the injected embryos, again demonstrating the presence of significant deletions in our samples.

### Optimisation of Cas9 and sgRNA concentration

Having demonstrated that we could visualise the presence of a large genomic deletion in our F0 samples, we next tested our ability to generate stable F1 lines (see below). At the same time, to improve the chance of successfully generating such mutants, we also focused on optimising the concentration of sgRNAs and Cas9 to be injected. As a positive control we targeted the pigmentation gene tyrosinase (*tyr*). We designed four different anti-*tyr* sgRNAs with target sites available in **figure S01**^13^. Although comparative studies across multiple genes and loci should be conducted to define optimal guidelines, we found that in our hands a concentration of 400pg of Cas9 associated with a molar ratio of 1:4 sgRNAs represents an attractive compromise between toxicity and total absence of pigment in the observed larvae; evidencing bi-allelic knockout, and thereby high efficiency of the cuts (**figure 2**&**S02**). Reducing the ratio of Cas9/sgRNA to 1:1 significantly reduced the efficiency as described below. We also compared yolk vs blastomeric injection which resulted in no clear difference in pigmentation loss. We therefore now perform our experiments with yolk-injection, facilitating the experimental procedure.

**Figure 02:**
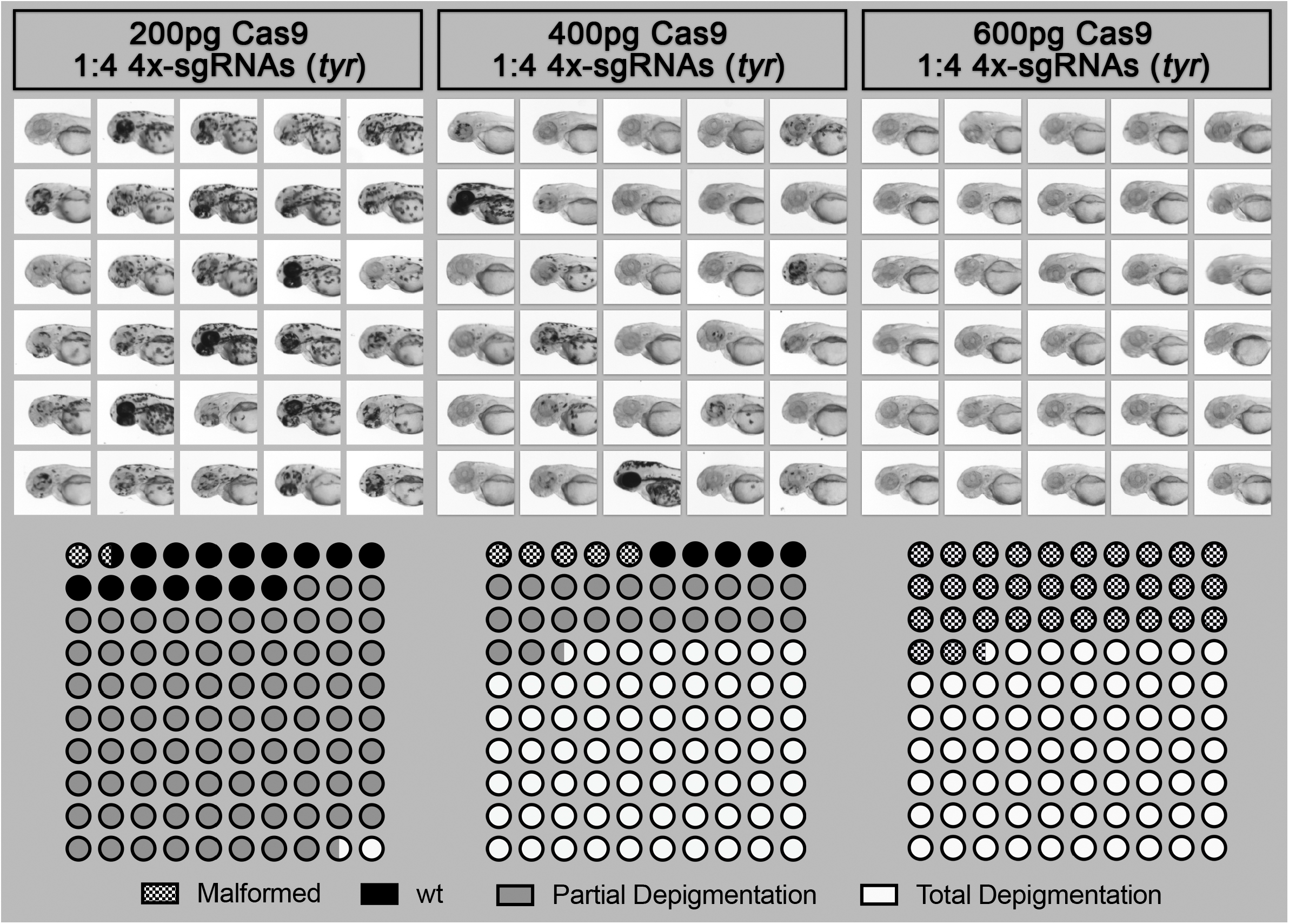
Effect of Cas9/sgRNA concentration on lethality, cutting efficiency and phenotype outcome in zebrafish. Different concentrations of Cas9 protein mixed with 4x sgRNA guides against the gene tyrosinase (*tyr*) were injected into the yolk of one-cell stage zebrafish embryos. Injected animals were then observed at 2dpf to score toxicity (malformation or death), partial depigmentation or total depigmentation (suggesting complete bi-allelic *tyr*-knockout). A mix of 400pg of Cas9/sgRNAs was selected to be used routinely in our pipeline.

### Pipeline for generating heritable stable deletions

To test our ability to generate stable large deletions across multiple loci, we worked with a mix of five different guides, four against our target and one against *tyr* (**tyr80, fig. S01, table S01**). We included an anti-*tyr* guide to help visualise correctly injected embryos and speed up the process of screening F0-founders (**fig. 1B**). At five days post-injection, we selected 80 injected larvae to grow based on their pigmentation phenotype, prioritising those with greater pigment loss (**fig. 1B and S02**). Once grown to adulthood, we outcrossed those with wildtype to determine if any F0 fish would transmit a stable large deletion in F1 embryos (**fig 1B**). As for the initial validation, it is straightforward to identify founders when the PCR reaction does not amplify a wt-amplicon as for total gene deletions. Therefore, only animals carrying such deletion would lead to a PCR product on the gel, justifying the pooling of embryos/DNA from a large population to evaluate potential transmission (**fig 1D**). However, for small deletions such as the transmembrane domain (≈0.15kb), amplification of the wt-target could mask the signal from potential indels, especially in cases of low transmission. Based on these results, for screens that do not involve the amplification of wt-amplicon, we routinely collect eight pools (seven with 10 embryos plus an extra mix of up to 50 embryos) per dish to further extract genomic DNA and screen for potential deletions/mutants by PCR (**fig 1D**). In contrast, for deletions involving the amplification of a wildtype band, we recommend not pooling the embryos, instead collecting seven individual embryos and only one pool of up to 10 embryos. Remarkably, we found at least one founder for each initially designed deletion. Importantly, we started our injections at a Cas9/sgRNAs molar ratio of 1:1 and obtained an average of 1 carrier/founder per 40 screened fish. Based on the results presented in **fig. 2**, we increased this ratio to 1:4 and observed an increase to approximately 1:10 founder. In total, we targeted the six *nrxn*-genes for the transmembrane domains (150kb deletion on average) as well as *nrxn1a* and *nrxn1b* for the entire gene (300kb and 70kb deletion respectively) and isoform-specific deletions (110kb and 22kb deletion respectively). Although we anticipated that the larger the deletion the harder it would be to isolate a mutant, this is not what we observed, with all deletions being obtained at a similar incidence. This contrasts with a recent study conducted by Wu *et al*., who found that the frequency of this type of deletion decreases with increasing distance between sgRNA target sites^14^. Finally, it is noteworthy that during the screens involving the deletion of the transmembrane domains, we found a high number of mutants presenting small indels at one or several sgRNA/target sites. Although the deletions or additions were relatively small, it was surprisingly easy to detect them on the gel facilitated by the formation of heteroduplexes. We were able to detect down to 2bp differences as a result of to the aberrant migration of the heteroduplex PCR products (**fig. 3**). This profile could be of great help to traditional approaches aiming to generatie small indels (frameshift mutants). Although it is unclear what promoted such formation/migration, we hypothesised that the size (300bp), as well as the position of the target sites (*i*.*e*. not in the middle of the amplicon), could play a role.

**Figure 03:**
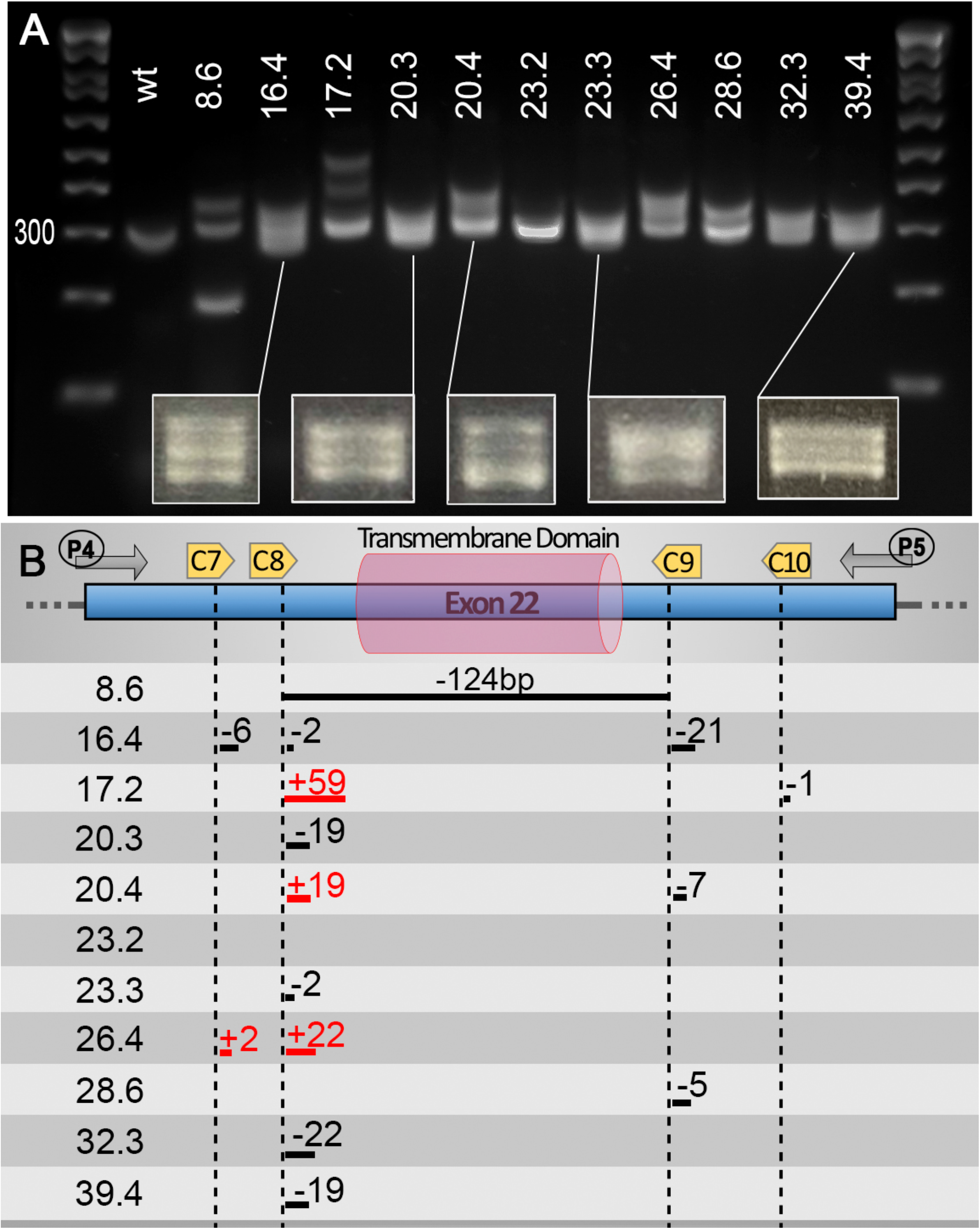
Heteroduplexes can help identify small indels using basic PCR screening approaches. Although not the primary aim of our pipeline, we found that, using a PCR screening approach, one can also easily detect “traditional” small indels due to an aberrant migration of heteroduplex PCR products. We found that at least a 2bp difference is sufficient to trigger a significant slower migration of potential heteroduplexes. We found that the heteroduplexes were best observed with an amplicon size of around 300bp, allowing clear visualisation of two or three bands on the gels. Additionally, most of our targets/cuts did not sit in the centre of the PCR amplicon, which might be a factor in promoting the formation of such heteroduplexes. **A**, Gel presenting migration of PCR amplicon using primers P4 and P5 on validated F1 animals (Number X.x, *i*.*e* 8.6, corresponds to fish number followed by embryo sample number). **B**, Mutations found in the different mutants are presented in the upper panel.

## CONCLUSION

Here, we demonstrated that large deletions ranging from hundreds of base pairs to hundreds of kilobases can be easily generated using multi sgRNA/target sites. Specifically, we found that the ratio of Cas9/sgRNAs has an effect on the founder rate whereas the size/length of the deletion does not. To generate stable mutants, a concentration of 400pg of Cas9 complexed with a molar ratio of 1/4 sgRNAs appears to offer an attractive approach, resulting in a final founder frequency of approximately 1 in 10 injected fish. **Supplemental table S03** provides a template for the design and preparation of such cocktails. It is worth noting that this ratio and concentration are supported by a recent study aiming at establishing a method for generating null phenotypes in G0 zebrafish^14^, although, for those “Morpholino-like” transient approaches, the authors found that 800pg of Cas9 complexed with a 1:6 molar ratio was best for generating F0 null mutants. However, one must consider that the potential toxicity and phenotypic consequences of bi-allelic mutations could potentially hamper the chance to generate stable lines. Our findings also demonstrate that the simultaneous injection of a “marker” sgRNA/target, such as an anti-*tyr* guide, can greatly facilitate the selection of properly injected F0 embryos as well as aiding the selection of F0 adults with a greater chance of mutation transmission in F1 generations. The downside is that one must counter-select F1 and F2 animals free of *tyr*-mutations. Finally, as presented in detail in the methods, we experienced problems generating high/sufficient yield for our guides using the various methods found in the literature. This led us to design an alternative approach based on full length oligonucleotides. This proved slightly more expensive, but yielded consistently large amount of good quality sgRNAs.

## AKNOWLEDGMENT

This work was supported by grants from the Australian National Health and Medical Research Council (NHMRC) Project Grant No 1165850 to BM and JG, The Rebecca Cooper Medical Research Project Grant No PG2019405 to JG. JG was also supported by a NHMRC Emerging Leader Fellowship No 1174145. We thank Rowan Tweedale for helping us review this manuscript.

## ONLINE METHODS

### Zebrafish Maintenance

Adult zebrafish and embryos were maintained by standard protocols approved by the University of Queensland Animal Ethics Committee. Ethic approval AE213_18/AE213_18. Wildtype lines used in these studies are TAB background.

### Genomic target selection and sgRNA Full length oligos design

Genomic sequences targeted by our single guide RNAs (sgRNAs) have been selected using both CRISPRscan (https://www.crisprscan.org/) and CHOPCHOP (https://chopchop.cbu.uib.no/) websites based on the zebrafish genome annotation (GRCZ11). Several methods are available online for producing sgRNAs to inject, however, we experienced two main problems: 1) inconsistent, low RNA yields with the template extension method and 2) long turnover times with the plasmid cloning/PCR method. This led us to optimise our own method resulting in the production of high yield and quality sgRNAs. Based on the target selected in either CRISPRSCAN or CHOPCHOP, we copied this region (with mutation if necessary) into a full-length oligonucleotide template (**supplemental figure/file S03**) which includes a T7 promoter in 5’ and the necessary Cas9 sequence in 3’. We then order sense (top) and antisense (bottom) full length oligos used in the “sgRNA preparation” presented below.

### sgRNA preparation and storage

Oligonucleotides (top and bottom oligos) were produced by Macrogen, Inc (South Korea, **Supplementary Table S01**) and reconstituted with Invitrogen Ultra-Pure Distilled water to 200µM. Double strand DNA templates for sgRNA synthesis were generated by annealing top and bottom oligos together with NEB buffer 2.1 in Thermal Cycler (Bio-Rad C1000 Touch) at 95°C for 5mins, with lid temperature at 105°C. Once the cycle was complete, tubes were left in Thermal Cycler, with lid closed, for 1 hour to allow slow temperature ramp down. The resulting template was purified using either Scientifix NucleoSpin Gel and PCR Clean-up Kit (#740609) or Invitrogen Gel Extraction and PCR Purification Kit (# K220001) according to manufacturer’s instructions, and eluted in 15uL Invitrogen Ultra-Pure Distilled water. We further quantified the purified DNA using Nanodrop (ThermoFisher ND-1000). Next sgRNAs were transcribed using a quarter reaction of the Ambion MEGAshortscript T7 Transcription Kit (#AM1354) with a DNA template concentration of 400-800ng and overnight incubation at 37°C. DNA templates were removed by addition of TURBO DNase at 37°C for 15mins. sgRNAs were then purified using Zymo Research RNA Clean&Concentrator Kit (R1015) and eluted in 15µL of Invitrogen Ultra-Pure Distilled water. Prior to storage at −80°C, we quantified the RNA using Nanodrop and checked RNA integrity on a 2% SB agarose gel.

### sgRNAs/Cas9 injection mix preparation

Each mix was prepared on ice the hour prior to injection. All solutions were transferred and prepared on ice. For calculating the amount to mix/pipette, we generated and used a template available in **supplemental table S03**. We used buffers and Cas9 protein from New England Biolabs (NEB, EnGen® Spy Cas9 NLS, # M0646). We prepared a 10µl mix that we stored on ice in a carrier box that was transported to the fish facility prior to setting up the injections.

### sgRNAs/Cas9 injections

One-cell stage or yolk injections were performed as previously described^15^. Male and female adult TAB wildtype were separated the day before injection. On injection-day, animals were mixed back together and monitored for mating behaviour. Animals were left mating for 30min after the release of the first eggs. In the meantime, injections were calibrated using a microscope micrometre calibration ruler. Embryos were then collected and distributed on a Sylgard injection tray. Depending on the experiment, we injected 1nl of Cas9/sgRNAs mix in either the yolk or the cell. Injected embryos were then collected and maintained in E3 medium supplemented with Methylene blue. Embryos were screened when necessary as described in the manuscript, and the selected larvae were transferred to the fish facility nursery for monitoring and feeding from 5-day post fertilisation (dpf).

### Genomic DNA extraction

Genomic DNA (gDNA) was extracted from embryos at 1dpf. Freshly prepared DNA extraction buffer (1M KCl, 0.5M Tris pH8, 1M EDTA, 20% IGEPAL, 10% Tween-20) with Proteinase K (10mg/mL) was added to embryo/s and digested in Thermal Cycler (Bio-Rad C1000 Touch) on the following program: 55°C for 2 hours and 98°C for 10 mins. gDNA was stored at −20°C. As described in the manuscript, for deletions involving the amplification of a wildtype band (transmembrane regions, figure 01), gDNA was extracted from individual zebrafish embryos (n=7) and a pool of ten embryos (n=1) for each F0/clutches screened. For deletions large enough to not involve the amplification of a wild-type band (whole gene or isoform-specific deletions), gDNA was extracted from 7x pool of 10 embryos and 1x pool of 50 embryos (whenever possible) for each F0/clutches screened.

### Validation via PCR Amplification

sgRNA/sCas9 cutting efficiency was evaluated by PCR amplification using primers specified in **figure 01** and **Supplemental Table S02**. We used AmpliTaq DNA polymerase (#N8080172) and followed manufacturer’s procedures. Briefly, PCR master mix was prepared with 10X PCR Buffer II, 25mM MgCl2, 10mM dNTP, 10uM forward and reverse primers, and AmpliTaq DNA Polymerase (5U/µL). 1µL of extracted DNA was added to the master mix in a 25µL final reaction volume and incubated in a Thermal Cycler. Conditions of the PCR amplification were: 95°C (1min), then 40 cycles at 95°C (30s) / 56°C (30s) / 72°C (20-30s), and a final extension at 72°C for 1min. PCR amplicons were revealed using gel electrophoresis using a 2% SB agarose.

### Sequencing&F1 generation

The amplified PCR products of samples identified as founders were sent for sequencing to Macrogen, Inc. (South Korea) using the same forward and reverse primers as the PCR screen. Sequencing results and mutations/deletions were processed using a combination of manual and automatic analysis. For the automatic analysis we used the following websites (http://yosttools.genetics.utah.edu/PolyPeakParser/; http://crispid.gbiomed.kuleuven.be/), however, in most cases those in silico approaches failed to help generate valuable information and did not lead to the identification of mutations. The best way to proceed was to manually read the Chromatograms and define the two alleles on a trial-and-error approach. Once zebrafish founders were confirmed with sequencing, these were out-crossed to wildtype TAB and grown for generating heterozygotes that would be identified through fin clipping.

